# Water column stability as an important factor controlling nitrite-dependent anaerobic methane oxidation in stratified lake basins

**DOI:** 10.1101/2021.11.09.467825

**Authors:** Guangyi Su, Moritz F. Lehmann, Jana Tischer, Yuki Weber, Jean-Claude Walser, Helge Niemann, Jakob Zopfi

## Abstract

Anaerobic oxidation of methane (AOM) with nitrate/nitrite as the terminal electron acceptor may play an important role in mitigating methane emissions from lacustrine environments to the atmosphere. We investigated AOM in the water column of two connected but hydrodynamically contrasting basins of a south-alpine lake in Switzerland (Lake Lugano). The North Basin is permanently stratified with year-round anoxic conditions below 120 m water depth, while the South Basin undergoes seasonal stratification with the development of bottom water anoxia during summer. We show that below the redoxcline of the North Basin a substantial fraction of methane was oxidized coupled to nitrite reduction by *Candidatus* Methylomirabilis. Incubation experiments with ^14^CH_4_ and concentrated biomass from showed at least 43-52%-enhanced AOM rates with added nitrate/nitrite as electron acceptor. Multiannual time series data on the population dynamics of *Candidatus* Methylomirabilis in the North Basin following an exceptional mixing event in 2005/2006 revealed their requirement for lasting stable low redox-conditions to establish. In the South Basin, on the other hand, we did not find molecular evidence for nitrite-dependent methane oxidizing bacteria. Our data suggest that here the dynamic mixing regime with fluctuating redox conditions is not conducive to the development of a stable population of relatively slow-growing *Candidatus* Methylomirabilis, despite a hydrochemical framework that seems more favorable for nitrite-dependent AOM than in the North Basin. We predict that the importance of N-dependent AOM in freshwater lakes will likely increase in future because of longer thermal stratification periods and reduced mixing caused by global warming.

## Introduction

Freshwater habitats such as lakes are important sources of methane (CH_4_), a potent greenhouse gas in the atmosphere (Bastviken et al. 2011). A large fraction of methane is produced in lake sediments by anaerobic methanogenic archaea, from where it may escape by ebullition or diffusion into bottom waters. Several studies have evidenced aerobic methane oxidation at the sediment surface or in the water column of lakes (He et al. 2012; Blees et al. 2014a; b; Milucka et al. 2015; Oswald et al. 2016). Within sediments or anoxic bottom waters methane may also be oxidized anaerobically. At least in the marine realm anaerobic oxidation of methane (AOM) is an important process mitigating methane emissions to the atmosphere (Knittel and Boetius 2009), and is mainly performed by microbial consortia of anaerobic methanotrophic archaea (ANME-1, -2 and -3) and sulfate-reducing bacteria (SRB) (Boetius et al. 2000; Michaelis et al. 2002; Orphan et al. 2002; Niemann et al. 2006). Recent studies have reported other potential electron acceptors for AOM, including nitrogenous compounds (Raghoebarsing et al. 2006; Ettwig et al. 2010; Haroon et al. 2013), iron and/or manganese (Beal et al. 2009; Sivan et al. 2011; Ettwig et al. 2016; Cai et al. 2018), and possibly humic substances (Scheller et al. 2016; Valenzuela et al. 2019).

Particularly in freshwater environments, AOM with electron acceptors other than sulfate may represent a significant methane sink (Sivan et al. 2011; Norði et al. 2013; Segarra et al. 2015; Weber et al. 2017; Su et al. 2020). For nitrogen-dependent anaerobic oxidation of methane (N-AOM), two different modes have been identified: The bacterial oxidation of methane with nitrite as terminal electron acceptor by *Candidatus* Methylomirabilis oxyfera (Ettwig et al. 2009; He et al. 2016; Versantvoort et al. 2018), where oxygen is produced intracellular disproportionation of nitric oxide to nitrogen and oxygen, which is then used for intra-aerobic methane oxidation (Ettwig et al. 2010). Secondly, true anaerobic oxidation of methane coupled to nitrate reduction, catalyzed by the methanotrophic archaeon *Candidatus* Methanoperedens nitroreducens (Haroon et al. 2013). Although the exact metabolic mechanisms of nitrate/nitrite-dependent AOM are not entirely elucidated, evidence for this process has been recently found in freshwater environments (Ettwig et al. 2009; Hu et al. 2009, 2014; Deutzmann and Schink 2011; Wang et al. 2012; Norði and Thamdrup 2014; Graf et al. 2018; Mayr et al. 2020) but also in marine oxygen minimum zones (Padilla et al. 2016).

Given the prevalence of nitrate in freshwater lakes, N-AOM may play an important role in the mitigation of methane emissions from lake sediments. In lacustrine environments, highest methane oxidation rates were often observed near oxic/anoxic transition zones at the sediment-water interface (Lidstrom and Somers 1984; Kuivila et al. 1988; Frenzel et al. 1990; Bender and Conrad 1994; He et al. 2012) or in the water column of stratified lakes (Rudd et al. 1974; Blees et al. 2014a, b). However, methane consumption at these boundaries was usually thought to be carried out by aerobic methanotrophs, fueled by oxygen supplied by diffusion, cryptic production, or intrusion events (Hanson and Hanson 1996; Bastviken et al. 2002; Pasche et al. 2011; He et al. 2012; Milucka et al. 2015; Oswald et al. 2016). Indeed, redox transition zones may also represent sites where nitrate/nitrite is typically produced/regenerated through the oxidation of ammonium and reduction of nitrate by nitrogen-transforming microorganisms (Kuypers et al. 2018). Hence, here the N-AOM might be masked by, or misinterpreted as, aerobic methane oxidation (Deutzmann et al. 2014). As a result, methane oxidation with nitrate/nitrite as terminal electron acceptor may play a greatly underappreciated role in lakes. While nitrate-dependent *Ca*. Methanoperedens has not been observed in a freshwater lake (Su et al. 2020), high abundance as well as transcriptional activity of nitrite-dependent *Ca*. Methylomirabilis limnetica has been recently reported in two permanently stratified lakes (Graf et al. 2018; Mayr et al. 2020). However, the ecology, and more importantly, the role of these denitrifying methane oxidizers may play in the lacustrine methane and nitrogen cycles still remain largely unknown.

In this study, we investigated methanotrophy in the anoxic waters of two main basins of Lake Lugano. Previous studies in this lake were mostly concerned with aerobic methane oxidation, and highlighted the prominent role of Type I methanotrophs near the redoxcline in the North Basin (Blees et al., 2014a), but also in the seasonal formation of a benthic nepheloid layer in the South Basin (Blees et al. 2014b). The data on the potential of AOM in the water column particularly of the North Basin remained ambiguous because potential methane oxidation rate maxima were found below the oxycline (Blees et al. 2014a). Here, we aimed at further elucidating the mode of methane oxidation (in particular the scope for true AOM), and the environmental factors that control the occurrence, growth, and activity of methanotrophs in the two contrasting lake basins. Lake Lugano is an excellent setting where to investigate the physico-chemical controls on AOM as a function of ecosystem dynamics, because the two hydrologically connected lake basins differ significantly in their mixing regimes, water column-stability, and hence redox conditions. We quantified methane oxidation rates in the water column of the two basins, performed incubation experiments to determine the effectiveness of different electron acceptors (nitrate, nitrite and sulfate) for AOM, and used 16S rRNA amplicon sequencing to identify key taxa of aerobic and anaerobic methanotrophic guilds. We demonstrate that *Ca*. Methylomirabilis is an important microbial player in anaerobic oxidation of methane in the North Basin. Moreover, archived DNA samples allowed us to track the population dynamics of *Ca*. Methylomirabilis and other methanotrophs in the years following an exceptional mixing event in 2005 and 2006.

## Materials and methods

### Site Description and Sampling

Lake Lugano is located at the Swiss-Italian border and consists of two hydrodynamically contrasting basins that are separated from each other by a shallow sill. The eutrophic 95m-deep South Basin undergoes seasonal stratification with the development of a benthic “bacterial” nepheloid layer and anoxia during summer and fall (Lehmann et al. 2004). The 288m-deep North Basin is permanently stratified, and a chemocline at about 100-130 m separates the oxic mixolimnion from the anoxic monimolimnion (Blees et al 2014a). The permanent stratification since the 1960’s was interrupted by exceptional mixing of the whole water column occured in 2005 and 2006 due to cold and windy winters, causing the transient oxygenation of the monimolimnion (Holzner et al. 2009; Lehmann et al. 2015).

Water samples were collected in late November 2016, in the center of the southern basin off Figino (45°57′N, 8°54′E), and off Gandria in the northern basin (46°06′N, 9°12′E). Oxygen concentrations were measured using a conductivity, temperature and depth (CTD) probe (Idronaut Ocean Seven 316 Plus). Water samples from distinct depths were collected using 5L-Niskin bottles, and subsamples were taken directly from the Niskin bottle and filtered (0.45 μm) and/or processed as outlined below. Water samples for methane oxidation potential measurements were collected in 20 mL glass vials, which were filled carefully through the tubing, allowing water to overflow for about 2-3 volumes. The bottles were filled completely and care was taken not to introduce any air bubbles. The vials where crimp-sealed with Br-butyl rubber stoppers (Niemann et al. 2015). Samples for methane concentration measurements were collected in 120 mL serum bottles, crimp sealed with thick butyl rubber stoppers (Niemann et al. 2015) and a 20 mL headspace was created before fixing the sample by adding 5 mL of 20% NaOH.

### Analytical Methods

Methane concentrations in the headspace of NaOH-fixed water samples were measured using a gas chromatograph (GC, Agilent 6890N) with a flame ionization detector and He as a carrier gas (Blees et al 2014b). Ammonium (NH_4_^+^) concentrations were determined colorimetrically using indophenol reaction, and nitrite (NO_2_^−^) using Griess reagent (Hansen and Koroleff (1999). NO_x_ (nitrate plus nitrite) was determined using a NO_x_-Analyzer (Antek Model 745). Nitrate (NO_3_^-^) concentrations were calculated from the difference between NO_x_ and NO_2_^−^. Filtered samples for sulfide (i.e. the sum of H_2_S, HS,^-^ and S^2-^) concentration determination were stabilized immediately after sampling with zinc acetate and analyzed in the laboratory photometrically (Cline 1969). Sulfate was analyzed by ion chromatography (881 IC compact plus pro, Metrohm, Switzerland). Water samples for dissolved iron (Fe^2+^) and manganese (Mn^2+^) were fixed with HCl (0.5 M final. conc) after filtration through a 0.45 μm membrane filter, and analyzed using inductively coupled plasma optical emission spectrometry (ICP-OES). Total Fe and Mn concentrations (unfiltered) were measured. Concentrations of Fe^2+^ were additionally determined photometrically using the ferrozine assay (Stookey 1970). Particulate iron was calculated from the difference between the total Fe^2+^ concentrations after reduction with hydroxylamine, and the dissolved Fe^2+^ in the filtered sample.

### Methane oxidation rate (MOR) measurements

In situ methane oxidation rates were determined with trace amounts of tritium-labeled methane (^3^H-CH_4_) (Steinle et al. 2015). Upon the retrieval of water samples, 5 μL anoxic ^3^H-CH_4_ solution (∼1.8 kBq) was injected into the 20 mL bubble-free glass vials and samples were incubated in the dark at 4 °C for 42 h. To terminate incubations, 2 mL water samples were directly transferred into 6 mL scintillation vials, mixed with 2 mL of a scintillation cocktail (Ultima Gold, PerkinElmer) and immediately measured for total radioactivity of ^3^H. 10 mL samples were then taken and transferred into 20 mL scintillation vials containing 1 mL saturated NaCl solution. After stripping the remaining radio-labeled methane from the vials for 30 min, samples were mixed with 8 mL of the scintillation cocktail prior to ^3^H_2_O radioactivity measurement via liquid scintillation counting (2200CA Tri-Carb Liquid Scintillation Analyzer). Methane oxidation rates (*MOR*) were calculated according to Eq. 1.

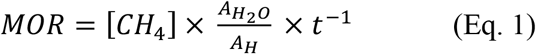

Where *A*_*H*_ and 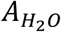represent the radioactivity of total ^3^H and ^3^H_2_O from methane oxidation, respectively, [CH_4_] is the methane concentration in the water column sample, and *t* the incubation time.

### Incubation experiments with ^14^C-labelled methane

We used microbial biomass from anoxic water layers of both basins to test different electron acceptors for their potential to stimulate anaerobic methane oxidation. Briefly, the biomass of a 500 mL water sample was collected on a glass fiber filter, which was then transferred to a 120 mL serum bottle containing 100 mL anoxic artificial lake water. The bottles were subsequently purged with nitrogen until the oxygen concentrations in the control bottles, equipped with trace oxygen sensor spots (TROXSP5, Pyroscience), were below the detection limit (0.1 μM). Under an N_2_-atmosphere in an anaerobic chamber, potential electron acceptors, i.e., nitrate, nitrite, and sulfate, were added from anoxic stock solutions to a final concentration of 4 mM, 4 mM, and 2 mM, respectively. Molybdate, a specific inhibitor of dissimilatory sulfate reduction, was added to some incubations (4 mM final concentration) to test for sulfate-dependent anaerobic methane oxidation. After these additions, the bottles were filled headspace-free with anoxic artificial medium, and closed with grey stoppers. The stoppers had been heated in boiled water, and were stored in a Schott bottle with Helium to remove dissolved oxygen in the elastomer. Finally, 10 μL of ^14^CH_4_ tracer were injected, and samples were incubated at 25 °C. To exclude potential oxygen contamination during the long incubation time, the closed incubation bottles were kept permanently under N_2_-atmosphere in an anaerobic chamber. Both live controls (without added electron acceptors) and base-killed controls (pH >13) were treated in the same way and incubated in parallel for the two basins. At the end of the incubation, 20 mL of headspace was created by exchanging the medium with N_2_ gas. Biological activity was stopped by adding 5 mL saturated NaOH solution (50% w/w). The radioactivity of residual ^14^CH_4_ (combusted to produce ^14^CO_2_), ^14^CO_2_ produced by methane oxidation, and radioactivity in the remaining samples was determined by liquid scintillation counting (e.g., Blees et al. 2014b). The first order rate constants (*k*) were calculated according to Eq. 2.

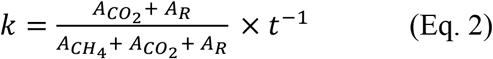

Where 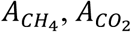, and *A*_*R*_ represent the radioactivity of methane, carbon dioxide, and the remaining radioactivity, respectively. *t* represents the incubation time. Methane oxidation rates (MOR) were calculated using the value for *k* and the methane concentration at the start of the incubation (Eq. 3).

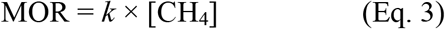

### DNA extraction, PCR amplification, Illumina sequencing and data analysis

Water samples from different depths of the two basins were collected and sterile-filtered for biomass using 0.2 μm polycarbonate membrane filters (Cyclopore, Whatman). Biomass DNA was then extracted using FastDNA SPIN Kit for soil (MP Biomedicals) following the manufacturer’s instructions. A two-step PCR approach (Monchamp et al. 2016) was applied in order to prepare the library for Illumina sequencing at the Genomics Facility Basel. Briefly, 10 ng of extracted DNA were used for a first PCR using universal primers 515F-Y (5′-GTGYCAGCMGCCGCGGTAA) and 926R (5′-CCGYCAATTYMTTTRAGTTT-3’) targeting the V4 and V5 regions of the 16S rRNA gene (Parada et al. 2016). The primers of the first PCR were composed of the target region and an Illumina Nextera XT specific adapter sequence. Four sets of forward and reverse primers, which contained 0-3 additional and ambiguous bases after adapter sequence, were used in order to introduce frame shifts to increase complexity (see Table S1 in the Supporting Information). Sample indices and Illumina adaptors were added in a second PCR of 8 cycles. Purified, indexed amplicons were finally pooled at equimolar concentration, denatured, spiked with 10% PhiX, and sequenced on an Illumina MiSeq platform using the 2×300 bp paired-end protocol (V3-Kit). After sequencing, quality of the raw reads was checked using FastQC (v 1.2.11; Babraham Bioinformatics). FLASH (Magoč and Salzberg 2011) was used to merge forward and reverse reads into amplicons of about 374 bp length with an average merging rate of 96%, allowing a minimum overlap of 15 nucleotides and a mismatch density of 0.25. Full-length primer regions were trimmed using USEARCH (v10.0.240), allowing a maximum of one mismatch. In a next step, the merged and primer-trimmed amplicons were quality-filtered (size range: 250-550, no ambiguous nucleotides, minimum average quality score of 20) using PRINSEQ (Schmieder and Edwards 2011). Clustering into operational taxonomic units (OTU) was done at a 97% identity threshold using the UPARSE-OTU algorithm in USEARCH v10.0.240 (Edgar 2010, 2013). Taxonomic assignment of OTUs was done using SINTAX (Edgar, 2016) and the SILVA 16S rRNA reference database v128 (Quast et al. 2013). Downstream sequence analysis was done in R v3.5.1 using Phyloseq v1.25.2 (McMurdie and Holmes 2013) as detailed in the supporting information (see supplementary method). Phylogenetic analysis of specific partial 16S rRNA gene sequences was performed in Mega 7 (Kumar et al. 2016) using the neighbor-joining method (Tamura et al. 2004), and the robustness of tree topology was tested by bootstraping (1000 replicates).

### Quantitative PCR (qPCR)

The abundance of *Ca*. Methylomirabilis was quantified using the primers qP1F (5′-GGGCTTGACATCCCACGAACCTG-3′) and qP1R (5′-CGCCTTCCTCCAGCTTGACGC-3′) amplifying positions 1001 to 1201 of 16S rRNA gene (Ettwig et al. 2009). qPCR reactions of all DNA samples were performed using the SensiFAST SYBR No-ROX Kit (Bioline) on a Mic (Magnetic Induction Cycler) real time PCR machine (BMS, Bio Molecular Systems, Australia). An initial denaturing step of 95 °C for 3 min was followed by 40 cycles of 5 s at 95 °C, 10 s at 65 °C, and 15 s at 72 °C. The specificity of the amplification was assessed by examining the melting curves from 60 °C to 95 °C, by agarose gel electrophoresis and sequencing. The calibration curves were generated using serial dilutions of pGEM-T Easy plasmid DNA (Promega, USA) carrying a single copy of the target gene fragment (qp1F/qp1R). Standard curves with these clones had a slope of -3.44, an R^2^ of 0.994, and an amplification efficiency of 95%. The number of gene copies in plasmid DNA was calculated using the equation reported previously (Ritalahti et al. 2006).

## Results and Discussion

### Hydrochemistry and methane oxidation in the water column of Lake Lugano

The water column of the deep North Basin (NB) was permantly stratified during the sampling year 2016 and oxygen concentrations decreased with depth and fell below the detection limit (1 μM) at 95 m depth. The redoxcline was defined between depth of oxygen starting to drop below 5 μM and depth of first occurrence of sulfide (Fig. 1A and S3). Methane concentrations increased linearly from redoxcline to 20 μM at 155 m, at depths where nitrite was below the limit of detection (0.02 μM) and nitrate were in the low micromolar range (< 1 μM, Fig. 1B). Below the redoxcline or in the anoxic water, concentrations of other reduced compounds such as sulfide, ammonium and dissolved Fe^2+^ rose above their background concentrations and increased continuously with water depth (Fig. S3). In the seasonally stratified South Basin (SB), the benthic nepheloid layer (NBL) stareted during summer and was fully developed in October. This turbid oxygen-depleted layer extends from the lake ground up to the chemocline, consists of microbial biomass, produced locally in large parts by methanotrophs (Blees et al. 2014b). During the sampling time in the southern basin, concentrations of oxygen decreased with depth and methane concentrations increased remarkably to 28 μM in the bottom water (Fig. 2A). Sulfide was below detection at all depths and the redoxcline was defined between depth at which oxygen drops below 5 μM and depth of first occurrence of reduced compounds such as dissolved Fe^2+^ (Fig. S4). The anoxic hypolimnia below redoxcline were characterised by considerable amounts of nitrate (38-70 μM), with low conccentrations of nitrite (1.2-3.9 μM) (Fig. 2B). Similar to the northern basin, ammonium accumulated in anoxic water column and increased with depth. Particular Fe reached up to 9.1 μM close to surface sediment and Mn species were present below the redoxcline (Fig. S4). Sulfate was relatively abundant and remained mostly above 100 μM below the redoxclines of both basins.

**Figure 1.**
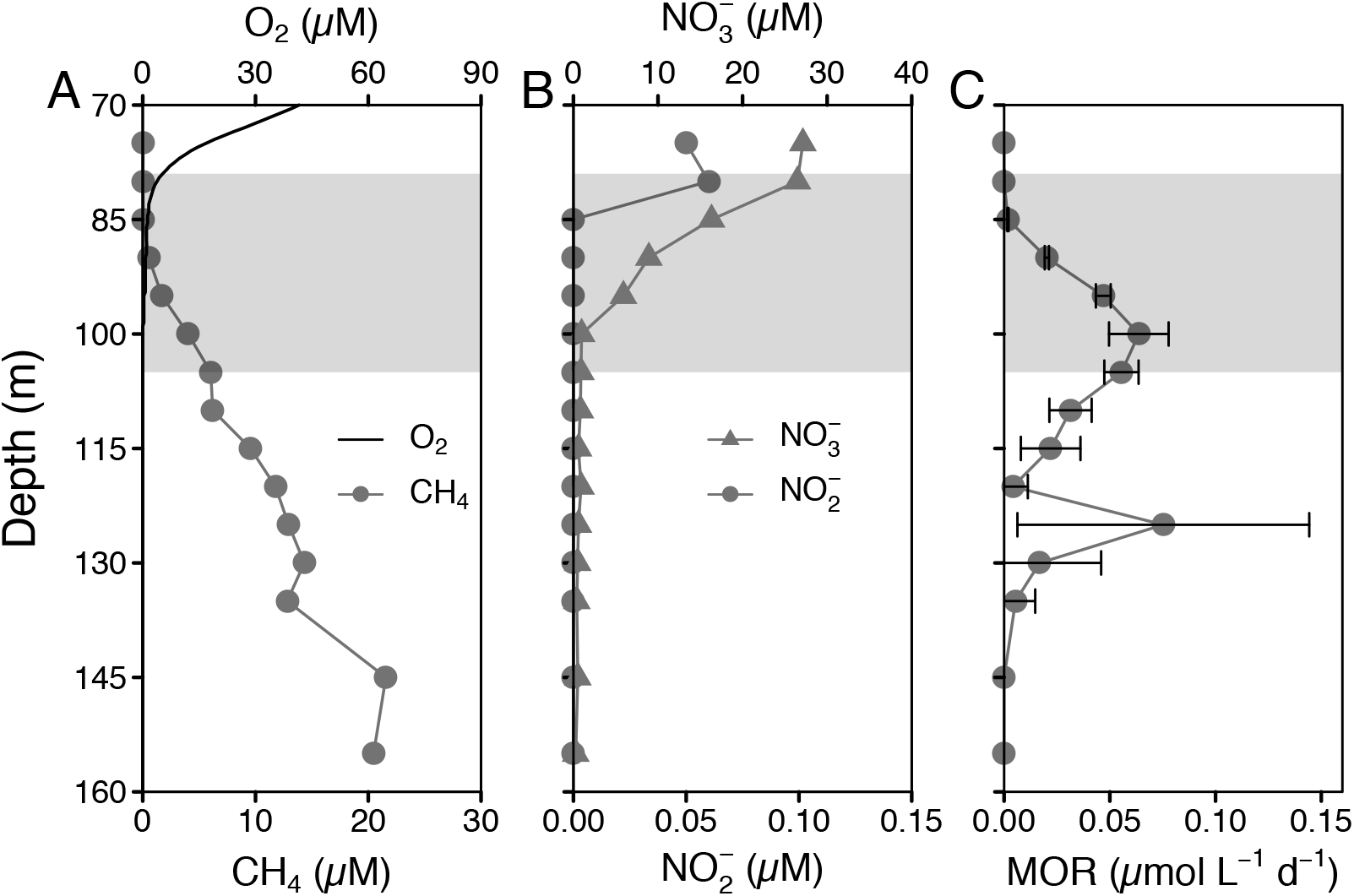
Water column profiles of (A) oxygen (O_2_) and methane (CH_4_) concentrations, (B) nitrate and nitrite concentrations, and (C) methane oxidation rates (MOR) in the North Basin of Lake Lugano in November 2016. The grey area represents the redox transition zone (RTZ), starting at O_2_ < 5 μM and reaching to the depth where sulfide rises above background concentrations. Error bars of MOR represent standard deviation (n = 3).

**Figure 2.**
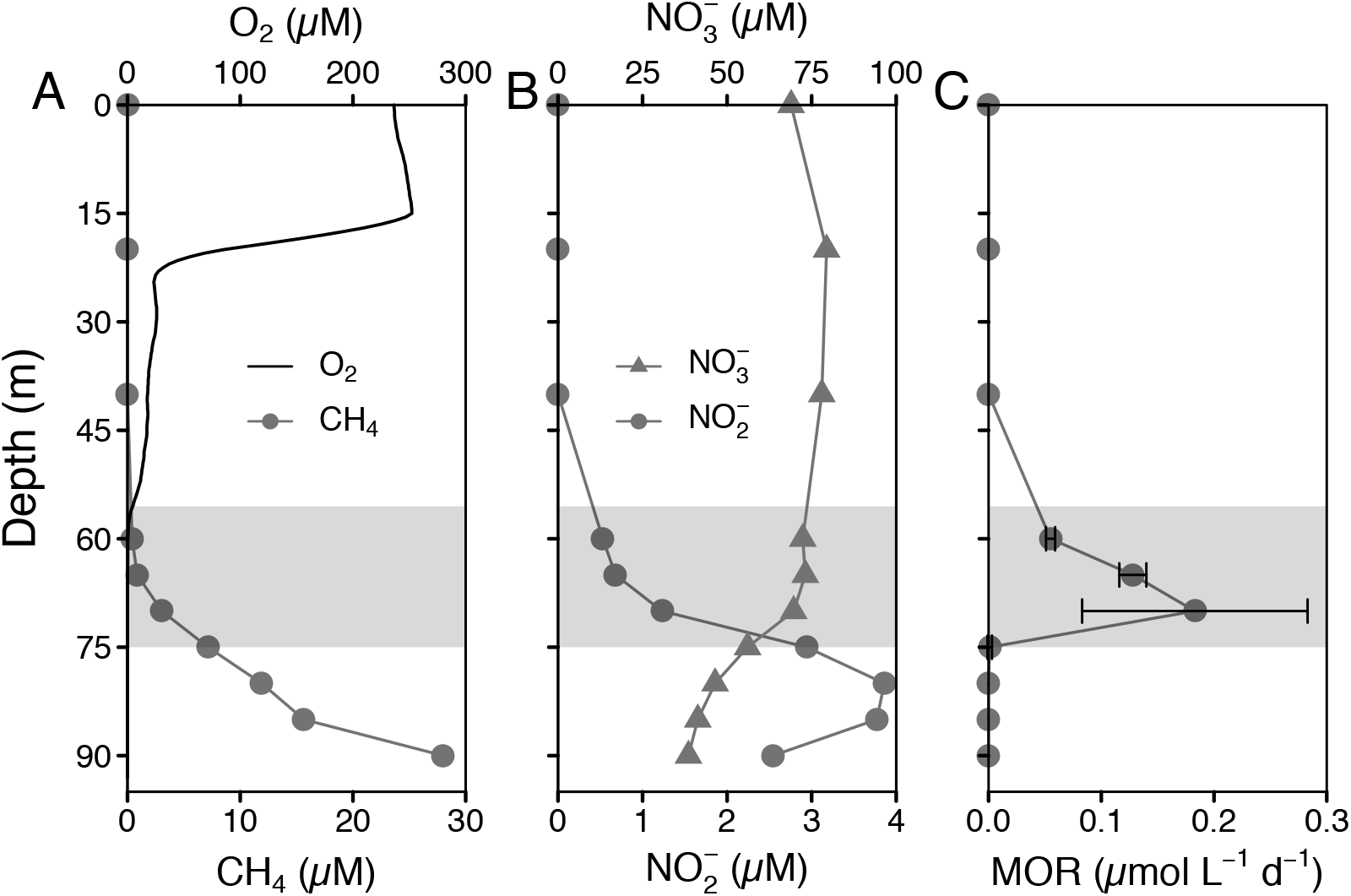
Water column profiles of (A) oxygen (O_2_) and methane (CH_4_) concentrations, (B) nitrate and nitrite concentrations and (C) methane oxidation rates (MOR) in the South Basin of Lake Lugano in November 2016. The grey area represents the RTZ. Error bars of MOR represent standard deviation (n = 3).

To quantify methane oxidation in the water column of Lake Lugano, we performed incubation experiments with ^3^H-CH_4_ to determine in situ rates of methane oxidation across the redoxclines particularly in the anoxic waters of both basins. In the NB, methane oxidation was occurring across and mostly below the redoxcline with two peaks observed (Fig. 1C). The first peak of methane oxidation (0.06 ± 0.01 μmol L^-1^ d^-1^) was at 100 m, right below the redoxcline, and aerobic methane oxidation was the most likely cause for this observed peak (Blees et al. 2014a). However, methane oxidation continued into the lower, anoxic parts of chemocline, and a secondary rate maximum of 0.08 ± 0.07 μmol L^-1^ d^-1^ was detected at 125 m. A similar bimodal pattern has also been observed before, albeit the two separate peaks and the oxycline were located at greater depths (Blees et al. 2014a). In contrast, we found a similar increase in methane oxidation rates across the redoxcline in the SB, with the highest rates of 0.18 ± 0.1 μmol L^-1^ d^-1^ observed within the redoxcline at 70 m (Fig. 2C). Although Type I methane-oxidizing bacteria (MOB) were shown to dominate the biomass in the BNL of SB, where the highest methanotrophic activity was observed (Blees et al. 2014a), it remained unclear whether the observed activity was soley due to these aerobic methanotrophs. The presence of both nitrate/nitrite and sulfate in the BNL bears the potential that methane could also be oxidized anaerobically with either of these oxidants.

Methane oxidation within oxic-anoxic transition zones of other meromictic lakes was often attributed to aerobic methanotrophs (Biderre-Petit et al. 2011; Oswald et al. 2016). In lakes with shallow redox transition zones (RTZs), cryptic oxygen production by phototrophs could sustain aerobic methane oxidation even in seemingly anoxic waters (Oswald et al. 2015; Milucka et al. 2015). At the depths of the RTZs in Lake Lugano, particularlay in the NB, oxygen production by phototrophs is an unlikely mechanism. Alternatively, Blees et al. (2014b) suggested that aerobic methane oxidizers can survive prolonged periods of oxygen starvation, and can resume high MOx activity upon episodic downwelling of oxygen, for example during cooling events. Yet potential mechanisms that inject oxygen to the deep hypolimnion were not investigated and it remains speculative if and how deep such events occur. Thus, methane oxidation far below the redoxcline in Lake Lugano North Basin may indeed be anaerobic, and nitrate/nitrite and sulfate may serve as potential oxidants for anaerobic methane oxidation.

### Evidence for nitrate/nitrite-dependent AOM

To test for the presence of active anaerobic methanotrophs, and to indentify the potential oxidants for methane oxidation, we set up anoxic incubation experiments with ^14^CH_4_ as substrate, different electron acceptors (i.e., nitrate, nitrite and sulfate), and concentrated biomass. The biomass was collected from 85-90 m in the South Basin, a depth well below the RTZ, but where nitrate, nitrite, and sulfate were present. Biomass from the North Basin was collected at 105-110 m, below the RTZ where nitrite was undetectable but nitrate (low) and sulfate were still available.

With biomass from water column right below the redoxcline of the meromictic NB, we found that both nitrate and nitrite stimulated AOM rates considerably (Fig. 3). Compared to the control experiments (i.e., no electron acceptor added), AOM rates increased by 52% (16 days, 54.8 ± 15.2 μmol L^-1^ d^-1^) and 72% (32 days, 60.8 ± 5.9 μmol L^-1^ d^-1^) in the presence of nitrate, and by 43% after 16 days (51.5 ± 6.3 μmol L^-1^ d^-1^) and 44% after 32 days (50.4 ± 15.8 μmol L^-1^ d^-1^) when nitrite was added. However, methane oxidation was not enhanced by the addition of sulfate compared to controls and there were no significant differences between the controls and amendments with sulfate of both the 16-day and 32-day incubations (Table S4). In addition, AOM rates were not significantly different between incubation bottles with sulfate and molybdate (Table S4). With respct to incubations with biomass from the anoxic water of the SB, no significant stimulation of methane oxidation was observed for all the treatments with added electron acceptors relative to the live controls after 16 days (Fig. 3 and Table S4). These results suggest that the oxidation of methane was not driven by any of the electron acceptors, and in turn, that AOM was likely not a major mode of methane removal in the South Basin, in spite of the presence of nitrate, nitrite and sulfate in the water column. Interestingly, after 32 days of incubation, the methane oxidation rates in all incubations were higher than after 16 days, independently of the added compounds, including molybdate, a known inhibitor of sulfate-reduction and thus of sulfate-dependent AOM (Wilson and Bandurski 1958).

**Figure 3.**
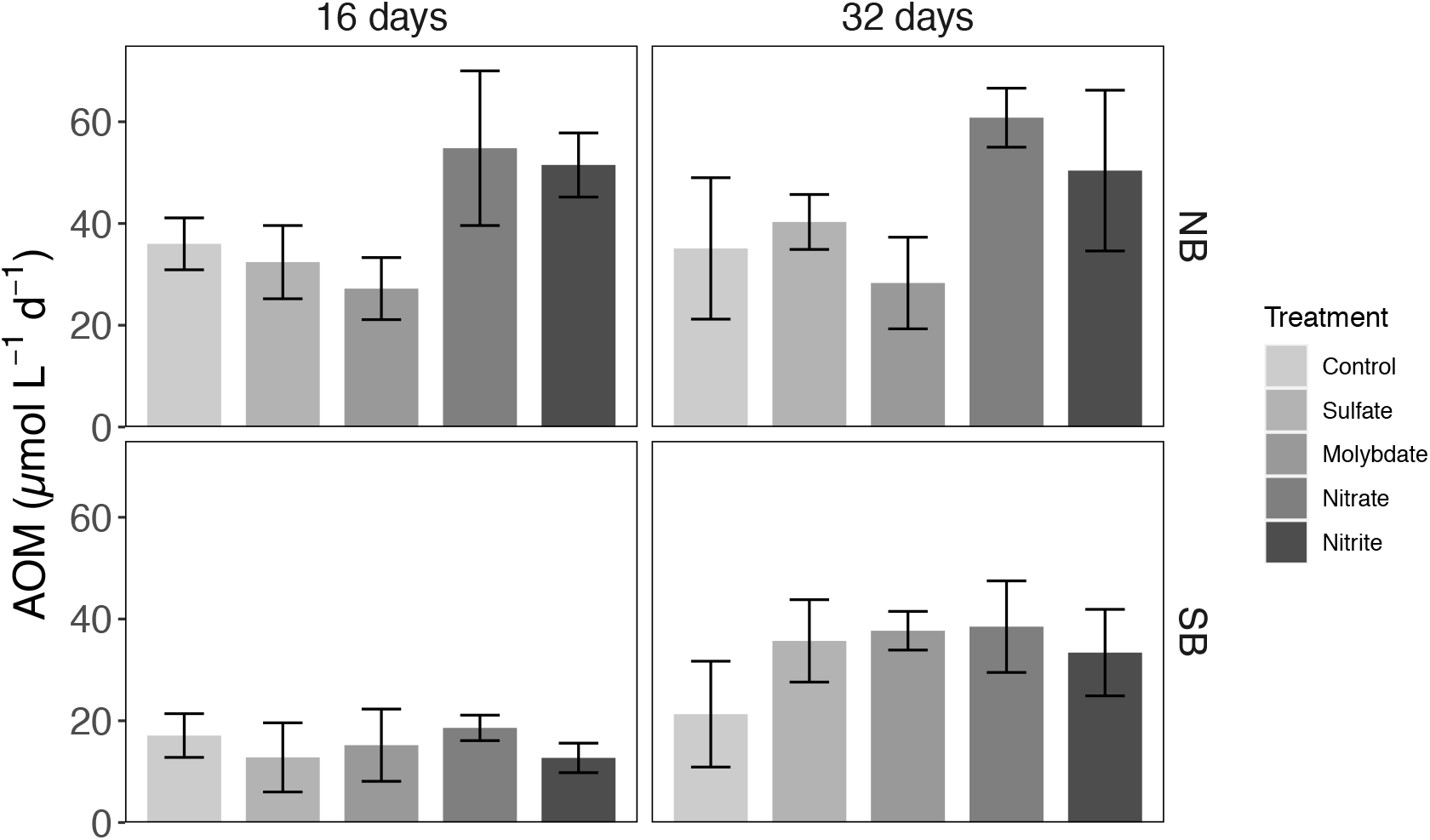
Effect of different electron acceptors on anaerobic methane oxidation (AOM) rates (n=3) in comparison to the control experiments (without addition of electron acceptors). The incubation was conducted with concentrated biomass collected in November 2016 from anoxic water layers in both NB and SB, and amended with ^14^CH_4_ and different oxidants (nitrate, nitrite, sulfate and molybdate). In the killed controls (n = 3), no tracer conversion was observed after 32 days.

Puzzling at first was the observation that we measured significant AOM rates in all live controls (both NB and SB), in the absence of any additional electron acceptors. Ambient concentrations of nitrate and nitrite were below the detection limit, but ambient sulfate was likely still present, which could explain the anaerobic oxidation of methane to some extent. However, the fact that no significant difference between sulfate- and molybdate-amended incubations was observed in all cases (Table S4) allows us to exclude any significant role of sulfate-dependent AOM in the live controls. One possiblity inferred from the incubation experiments particularly in the SB was that the methanotrophs in the concentrated biomass oxidized methane with the particulate Fe and/or Mn oxides (Beal et al. 2009; Ettwig et al. 2016; Cai et al. 2018), which were present in the water column and were concentrated on the filters together with the biomass (Fig. S3 and S4). In addition, the presence of traces of oxygen cannot be excluded. Greatest care was taken to avoid any oxygen contamination during preparation of the incubations and sampling. We did not, however, add a reducing agent to remove chemically any traces of oxygen. Thus, if trace amounts of oxygen (<100 nM) were still present in the incubations, they might be sufficient to serve as substrate for the enzyme methane monooxygenase. After the oxygen-dependent initial attack of methane, its further transformation could proceed anaerobically by fermentation, whereby hydrogen, formate acetate, and other compounds are produced (Kalyuzhnaya et al. 2013). Because short chain fatty acids are volatile under acidic conditions, radiolabeled formate and acetate could have been purged from the samples together with CO_2_, and contribute to the measured methane oxidation rate. This fermentative conversion of methane to excreted organic compounds by gammaproteobacterial methanotrophs, could represent an important methane elimination pathway under severe oxygen limitation (Kalyuzhnaya et al. 2013) in both the nepheloid layer of the SB and water column below the RTZ of the NB of Lake Lugano.

Although the electron acceptors used for methane oxidation in all live controls remained to be explored, incubation experiments with biomass from the NB demonstrated that both nitrate and nitrite played a significant role and enhanced methane oxidation under anoxic condition, providing evidence for methane oxidation coupled to nitrate and/or nitrite reduction. Assuming that methane consumption in the live controls was an effect due to micro-aerobic methane oxidation or metal-dependent AOM, rates in the nitrite/nitrate amendments beyond the controls (net consumption) can be attributed to true NO_x_-dependent AOM (N-AOM). Indeed, nitrate/nitrite-dependent AOM contributed 52-73% and 43% to the overall methane consumption in nitrate/nitrite-added incubations with water from below the chemocline in Lake Lugano North Basin, hightlighting the potentially large role N-AOM plays in lacustrine methane cycling.

### Abundance of diversity of methanotrophic bacteria

Based on 16S rRNA amplicon sequencing using an optimized PCR cycle (Supplementary method), a total of 4518 OTU’s were obtained in the combined datasets of the South and the North Basin, with a total of 7247 OTU’s before rarefaction (Weiss et al. 2017). Among them, we identified 32 OTU’s of potential methanotrophic bacteria, 23 of which were related to gamma-proteobacterial (Type I) methanotrophs. One OTU was related to Methylomirabilia, the former NC10 class (Fig. S5). No evidence of typical anaerobic methane-oxidizing archaea such as *Ca*. Methanoperedens or representatives of the ANME-1, -2 or -3 groups were detected in any of the samples from the water column of both basins.

In the permanently stratified NB, we detected 16S rRNA gene sequences that were affiliated with *Ca*. Methylomirabilis, which was capable of mediating AOM with nitrite as terminal electron acceptor (Ettwig et al. 2010). With a relative sequence abundance of up to 6.7% at 95 m, this single OTU was equally abundant as all *Methylococcaceae* sequences combined, which accounted for ∼5% of total sequences within or below the redoxcline (Fig. 4 and 5, Nov. 2016). Both *Ca*. Methylomirabilis and *Methylococcaceae* coexisted in the micro-oxic water column (85-105 m), where methane consumption rates were high. These observations together with incubation experiment from the NB suggested that both aerobic and anaerobic methanotrophs were important members of the methanotrophic guild and contributed together to the efficiency of the pelagic methane filter close to the redoxcline (Fig. 1). However, the factors facilitating the secondary peak of methane oxidation activity at 125 m (i.e., in anoxic water) remain uncertain. The lack of evidence for both sulfate or iron-utilizing anaerobic methanotrophs suggested that this peak was most probably attributed to *Ca*. Methylomirabilis (Fig. S6). Although the abundance of this denitrifying methane oxidizer did not vary in tandem with AOM rate at that depth (1.5% rel. abundance) and nitrite concentrations were below the detection limit throughout the water column, *Ca*. Methylomirabilis may still depend directly on the in-situ production of nitrite through microaerobic ammonium oxidation or nitrate reduction. Indeed, relatively high concentrations of ammonium were observed at these depths, and the abundance of ammonium-oxidizing bacteria (AOB) was peaking at 130 m (0.42% relative abundance), closed to the depth where the secondary methane oxidation maximum was observed (Fig. S7B). Oxygen was not detected at this depth, but these microorganisms could utilize minimal oxygen concentrations (at low nanomolar levels), which occurred at redoxcline boundaries or through sporadic oxygen intrusions oxygen and sustained aerobic ammonium oxidation (Thamdrup et al. 2012; Bristow et al. 2016). Thus, the virtual absence of nitrite was a reflection of the very efficient consuption rather than a lack of production, which implies that well below the chemocline of the NB, the nitrite-dependent AOM was likely limited by the nitrite production processes.

**Figure 4.**
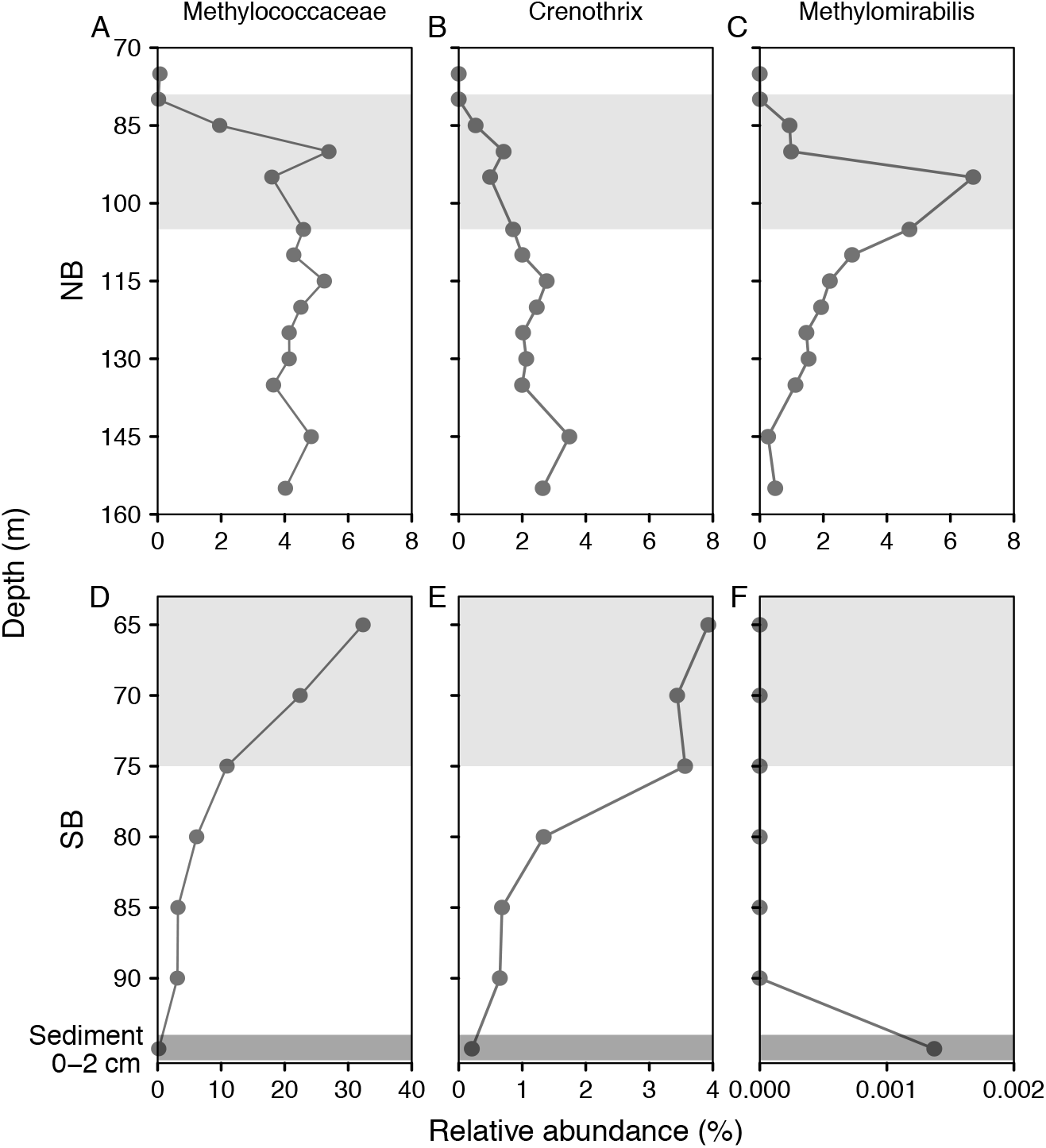
Relative abundance (% of total sequences) of (A) *Methylococcaceae*, (B) *Crenothrix* and (C) *Ca*. Methylomirabilis in the water column of Lake Lugano South Basin in November 2016. Data are based on relative read abundances of 16S rRNA gene sequences. The RTZ (light grey) or surface sediment (0-2 cm, dark grey) is represented by the grey shaded area.

**Figure 5.**
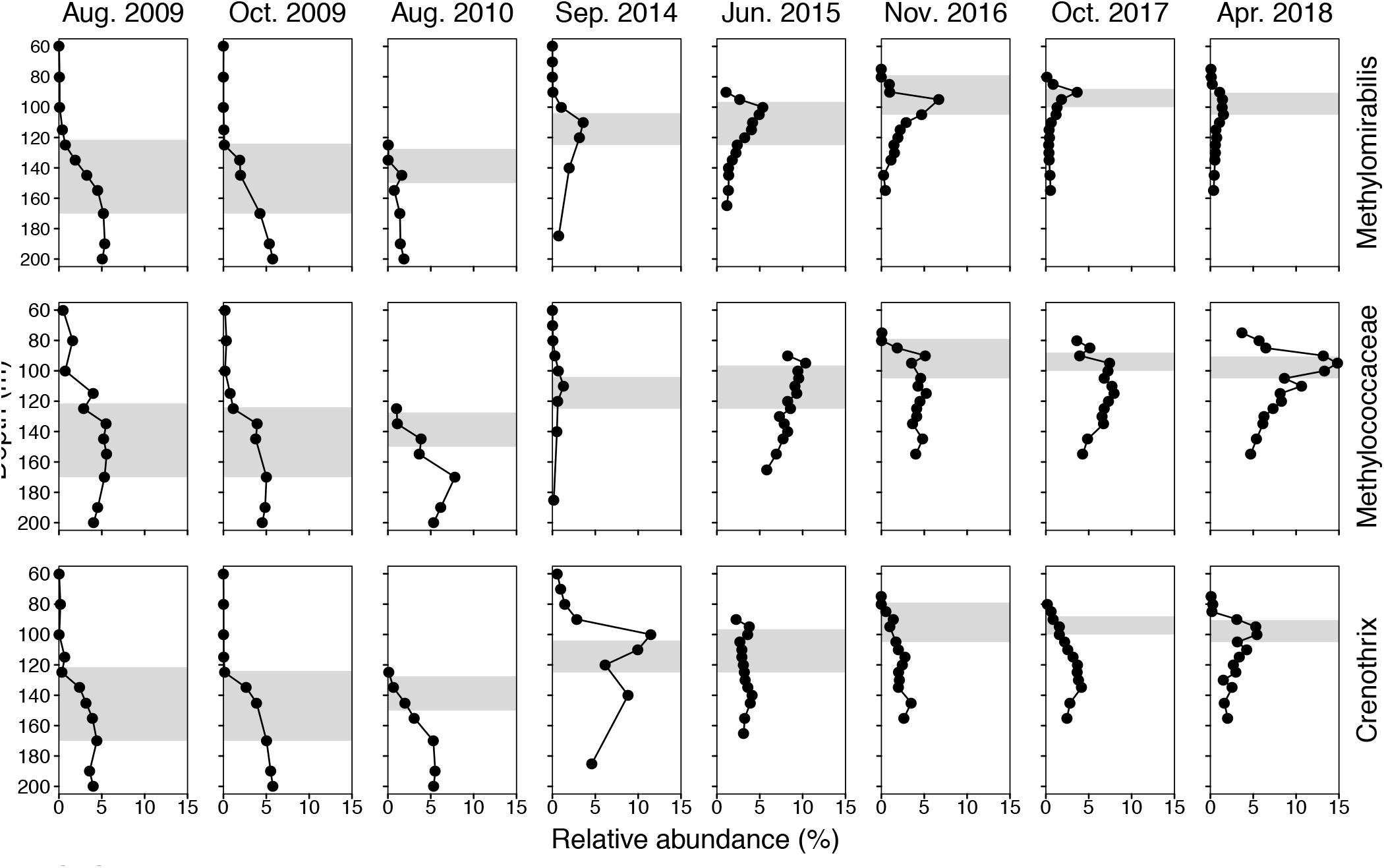
Time series data of *Ca*. Methylomirabilis, *Crenothrix* sp, and *Methylococcaceae* in the water column of Lake Lugano North Basin, starting three years after the exceptional mixing events in 2005/2006. Data are based on relative read abundances of 16S rRNA gene sequences. Grey areas represent the location and extension of the RTZ for the different sampling dates.

By comparison, we were not able to detect any 16S rRNA gene sequences that belonged to *Ca*. Methylomirabilis in the water column of the seanonally stratified SB, but only in the surface sediment with a very low relative abundance of 0.0014% of total sequences (Fig. 4F). The apparent absence of true anaerobic methane oxidizers was consistent with the lack of any significant stimulation of AOM with nitrate/nitrite in the incubations with biomass from this basin. However, Type I methane-oxidizing bacteria or *Methylococcaceae* were dominating the aerobic methanotrophic community (Fig. 4D), which was in accordance with the previous finding (Blees et al. 2014b). Among the members of *Methylococcaceae* (including *Methylobacter, Methylomonas, Methyloglobulus, Methylosarcina, Methyloparacoccus, Methylocaldum* and *Ca. Methylospira*), *Methylobacter* represented the most abundant genera, followed by *Crenothrix* (Fig. 4B), which has recently been shown to contribute to methane oxidation in two other stratified lakes in Switzerland (Oswald et al. 2017). Despite their occurrence in anoxic waters, these gamma-protebacterial methanotrophs are considered aerobic methanotrophs, as they use molecular oxygen for the particulate methane monooxygenase and the initial activation of methane. Nevertheless, genomes of several aerobic methanotrophs, including *Crenothrix*, encode putative nitrate (*narG, napA*), nitrite (*nirS, nirK*), and/or nitrogen oxide reductases (*norB*) (e.g. Kits et al. 2015; Oswald et al. 2017). *Methylomonas denitrificans*, for example, can couple the oxidation of methane (and methanol) to the reduction of nitrate to N_2_O, under severe oxygen limitation (Kits et al. 2015). But oxidation of methane under completely anoxic conditions has neither been shown for *Crenothrix* nor any other of the gamma-proteobacterial methanotrophs to date. When comparing the diversity of *Methylococcaceae* in the water column of the two basins, six out of the total 23 OTU’s were shared by both basins, 4 OTU’s were observed in the North Basin only, and 13 in the South Basin only (Fig. S5). Thus, the methanotrophic guild was more diverse (19 OTUs) in the dynamic water column of the SB, where oxic and anoxic conditions alternate seasonally.

Taken together, the peak activity of methane oxidation at 70 m in the SB was essentially due to aerobic *Methylococcaceae* (22% rel. abundance) thriving under hypoxic conditions in the BNL. More importantly, in the NB of Lake Lugano, methane oxidation within the redoxcline was mediated by both type I MOB and nitrite-dependent *Ca*. Methylomirabilis. Furthermore, we propose that the observed methane oxidation in the seemingly anoxic water column was largely attributed to denitrifying methanotrophs using cryptic nitrite from “nano-aerobic” bacterial nitrification, but we could not exclude the cryptic aerobic methane oxidation. Independent of the mode of methane oxidation (aerobic or anaerobic), oxygen appears as the ultimate and key limiting factor that controls methane oxidation directly (MOx) or indirectly (nitrite-dependent AOM) in the North Basin. Yet, it remains enigmatic as to why we see the accumulation of aerobic and nitrite-dependent methanotrophs, and even more intriguingly a methane turnover rate peak, at a very specific water depth well below the redoxcline (Fig. 1C).

### Water column stability as an ecological factor fostering nitrite-dependent anaerobic methane oxidation

After ∼40 years of permanent meromixis in the North Basin, two exceptionally strong mixing events in 2005 and 2006 led to a complete oxygenation of the entire water column (Holzner et al. 2009; Lehmann et al. 2015). Thereafter, the water column re-stabilized rapidly again and remained stratified with anoxia below 125 m depth (e.g., Wenk et al. 2013). Interestingly, in 2009 the abundance of *Ca*. Methylomirabilis was low around the redoxcline but increased with depth reaching ∼5% of total sequences at 200 m. The vertical mixing brought considerable amounts of oxygen to the deep water column and resulted in the almost complete oxidation of ammonium, the produced nitrite and nitrate fueled the growth of *Ca*. Methylomirabilis. Once nitrite was used up, the abundance of this methanotrophic bacteria started to decrease in the monimolimnion. On the other hand, a new population of Ca. Methylomirabilis started to establish at the RTZ (Fig. 5). Notably, an upward migration of the redoxcline occurred after 2010 but the peak abundances of *Ca*. Methylomirabilis were consistently observed within the redoxclines at 110 m, 100 m and 95 m, respectively, representing up to 3.6% of total sequences in September 2014, 5.4 % in June 2015 and 6.7% in November 2016. The steadily increased relative abundances of *Ca*. Methylomirabilis based on read numbers were further confirmed by qPCR (Fig. S8), demonstrating that water column stability was an important environmental factor for the growth of nitrite-dependent anaerobic methanotrophs. Further putative evidence is provided when comparing the two lake basins. The chemical conditions in the deep water column of the SB were conducive to nitrate/nitrite-dependent AOM (e.g., nitrate and nitrite concentrations up to 73 and 3.9 μmol/L, respectively, Fig. S4), but *Ca*. Methylomirabilis was not detected at all where nitrite-dependent AOM should be operative and was only found in the anoxic surface sediment. These observations suggest that the seasonal mixing regime with a stratified anoxic period of ∼5 months (shorter than our estimated doubling time of ∼6 months, Fig. S9) did not support the development of stable populations of the slow-growing anaerobic methanotrophs and stable water column condition was a more critical factor than previously thought. In addition, ventilation of hypolimnetic waters in 2017 and 2018 resulted in a decline of the redoxcline and the strikingly decrease of the abundance of *Ca*. Methylomirabilis (Fig. 5). However, this oxygenation did not affect but appeared to stimulate the growth of *Methylococcaceae* above the redoxcline, indicating that they were most likely outcompeting the denitrify anaerobic methanotrophs that migh be intolerant of high oxygen concentrations.

Based on the existing time-series data demonstrating the increasing relative importance of *Ca*. Methylomirabilis in the North Basin, we speculate that the observed evolution reflects the slow dynamic recovery of the nitrite-dependent AOM community after the mixing events in 2005/2006. Episodic mixing and ventilation of hypolimnetic waters marks a significant ecological perturbation, which likely has detrimental effects particularly on this slow-growing anaerobic methanotrophic bacteria that require stable and low-redox environmental conditions (Luesken et al. 2012). Once quasi-permanent anoxia under stably stratified conditions was restored, the population *Ca*. Methylomirabilis seemed to grow back in the deeper hypolimnion, and remained a permanent and important component of the water-column methane filter in the North Basin.

## Conclusions

In this study, we have shown that nitrite-dependent AOM was an important methane sink in the permanently stratified North Basin of Lake Lugano and that this process was mediated by *Candidatus* Methylomirabilis below the redoxcline and in the anoxic water column. Time series data demonstrate that stable and low redox conditions in the meromictic North Basin are particularly conducive to the development of AOM-performing bacterial methanotrophs. In the more dynamic South Basin, the duration of seasonal stratification and anoxia is likely too short, relative to the slow growth rate of *Ca*. Methylomirabilis, to allow the establishement of a stable population, in spite of the favorable hydrochemical conditions with high dissolved nitrate/nitrite and methane concentrations. Our research on methanotrophy in the two connected but hydrodynamically differing lake basins highlights that the chemical conditions alone cannot fully determine which microorganims will thrive and prevail in a system. Instead, physical processes such as water column dynamics/stability may be equally important, but often neglected, factors determining microbial community structures, and with that the modes and activity of biogeochemical processes. Our findings link water column stability and nitrite-dependent AOM within the stratified lake and have important implications for both anaerobic processes and the prediction of future methane emission under the scenarios of climate change.

## Supporting Information

### Supplementary Methods

#### Library preparation

The amplification conditions of the initial PCR during library generation are critical and can significantly affect results. Too few amplification cycles will limit the detection of low abundant taxa, too many will lead to PCR product saturation and to a biased representation of microbial community structures. In order to optimize our library preparation protocol, we evaluated the effect of two different amplification cycle numbers (18 vs. 25 cycles) for the initial PCR step, resulting in two libraries, which were sequenced in the same Illumina run.

The initial PCR consisted of 12.5 μl of 2X KAPA HiFi Hot Start Ready Mix, 0.75 μl forward primer (10 μM), 0.75 μl reverse primer (10 μM), 1 μl of PCR grade water, and 10 μl of template DNA (1 ng/μl). Four sets of 16S rRNA primers (Table S1) were used to amplify sample DNA. The primers contained 0-3 additional ambiguous bases between the adapter sequence and the PCR primer 515F-Y/926R (Parada et al. 2016) to increase the nucleotide diversity and improve template generation during Illumina sequencing. Increasing the nucleotide diversity at this stage allowed using a relatively low amount of PhiX (10%) during sequencing. PCR was run on a Biometra thermocycler using the following program: 98 ºC for 3 min; cycles of 98 ºC for 20 seconds, 55 ºC for 15 seconds, and 72 ºC for 15 seconds (18 or 25 cycles, respectively); 72 ºC for 5 minutes final elongation. Samples were then cleaned using AMPure Beads following the manufacturer’s instructions. Nextera XT index primers (N7XX and S5XX; Illumina) were attached to the amplicons in a subsequent 25 μl PCR reaction with: 12.5 μl of 2X KAPA HiFi, 2.5 μl each of Nextera XT index primer 1 and 2, 2.5 μL of PCR water and 5 μl of the cleaned amplicon run at 95 ºC for 3 min; 8 cycles of 95 ºC for 30 seconds, 55 ºC for 30 seconds, and 72 ºC for 30 seconds; 72 ºC for 5 minutes. Products were again cleaned using AMPure Beads and quality checked using Agilent Fragment analyzer (dsDNA-915 reagent kit), quantified using Qubit (ThermoFisher), normalized and finally pooled at equimolar quantities. Sequencing was done on a MiSeq system (Illumina) using the PE 300 method (V3 reagent kit) with 10% PhiX addition. Library preparation and sequencing was done at the Genomics Facility Basel (D-BSSE ETHZ and Basel University). Raw sequence data are made available at NCBI under the BioProjectID PRJNA672280 with the accession numbers from SRR12936362 to SRR12936382.

Quality control of raw reads, initial sequence treatment, and taxonomic assignment is outlined in detail in the main manuscript. Sequencing results are summarized in Table S2. 16S rRNA sequence data were analyzed in R (v3.5.1) (R Core Team, 2014, http://www.r-project.org/) using mostly the libraries: phyloseq (McMurdie and Holmes 2013), vegan (Oksanen et al. 2013), ggplot2 (Wickham 2009). Rarefaction curves show that 25 cycles yielded better estimates of the species richness than 18 cycles (Figure S1). ANOVA in R was used to test whether different alpha diversity measures (i.e., Observed richness, Chao1, Shannon and InvSimpson) were affected by either the sequencing depth or the amplification cycle number. Results show (Table S3) that Shannon and InvSimpson are significantly affacted by the amplification cycle number, while others (e.g., Observed richness and Chao1) depend on the sampling depth (Weiss et al. 2017). Accordingly, samples were rarefied for alpha diversity estimates, or where the numbers operational taxonomical units (Z)OTUs in the two libraries were compared (Table S2). Principal coordinate analysis (PCoA) was performed after data were rarefied to the same depth. PCoA plots show that both 18 or 25 PCR amplification cycles yield very similar microbial community structure (Figure S3). Ultimately we choose 25 amplification cycles for the microbial community analysis, because of the higher sensitivity minor taxa. The sample G_100m in the 25 cycle library was characterized by particularly low read and (Z)OTU numbers (Table S3), as well as an atypical community structure (Figure S2), suggesting that it was affected by an amplification artifact. It was therefore excluded from any further analysis.

## Supplementary tables

**Table S1.**
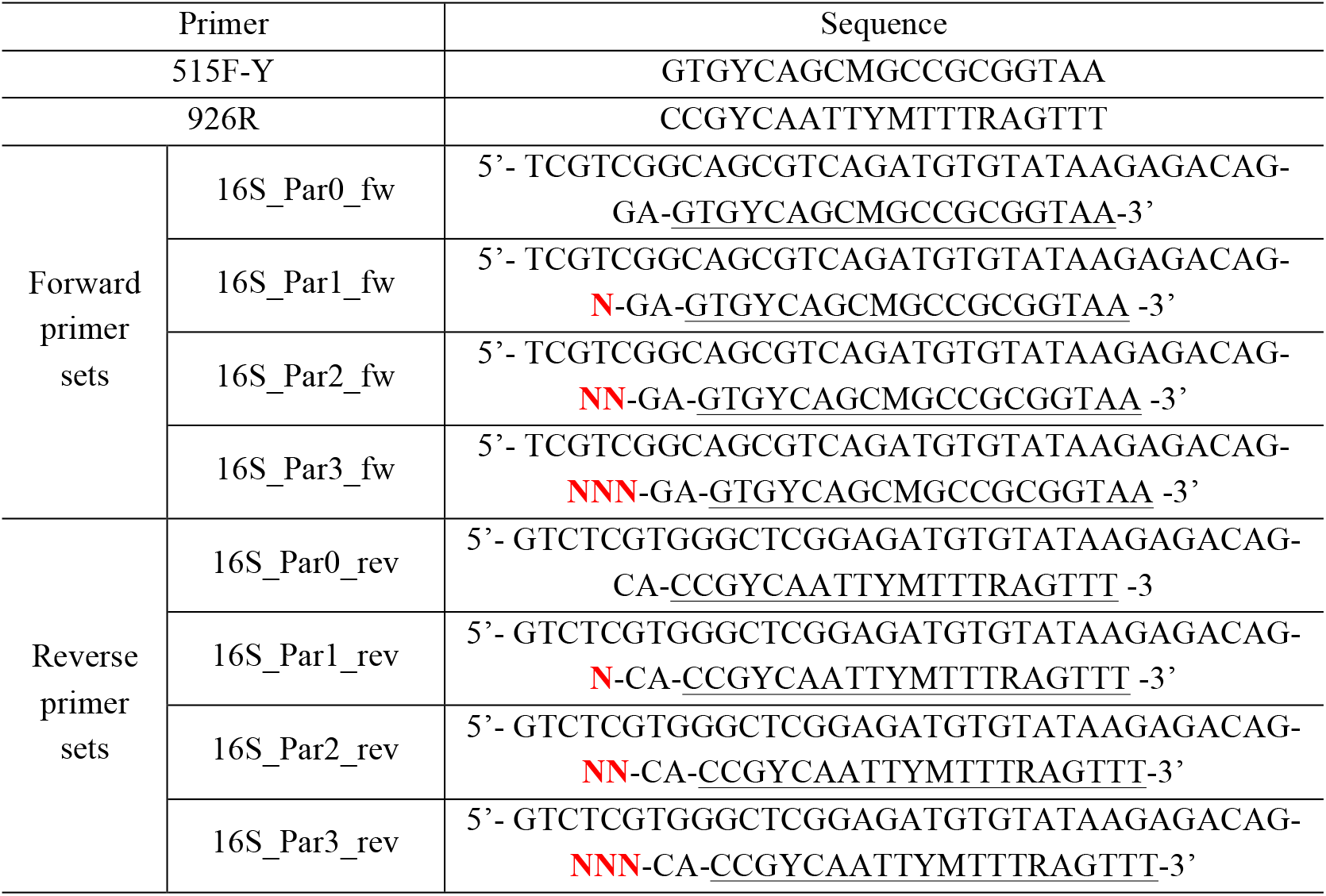
Design of the four different sets of forward and reverse primers used for library preparation. The primers contained 0-3 additional ambiguous bases (indicated in bold red) between the adapter sequence and the amplicon PCR primer 515F-Y/926R (Parada et al. 2016) to increase the nucleotide diversity and improve template generation during Illumina sequencing.

**Table S2.**
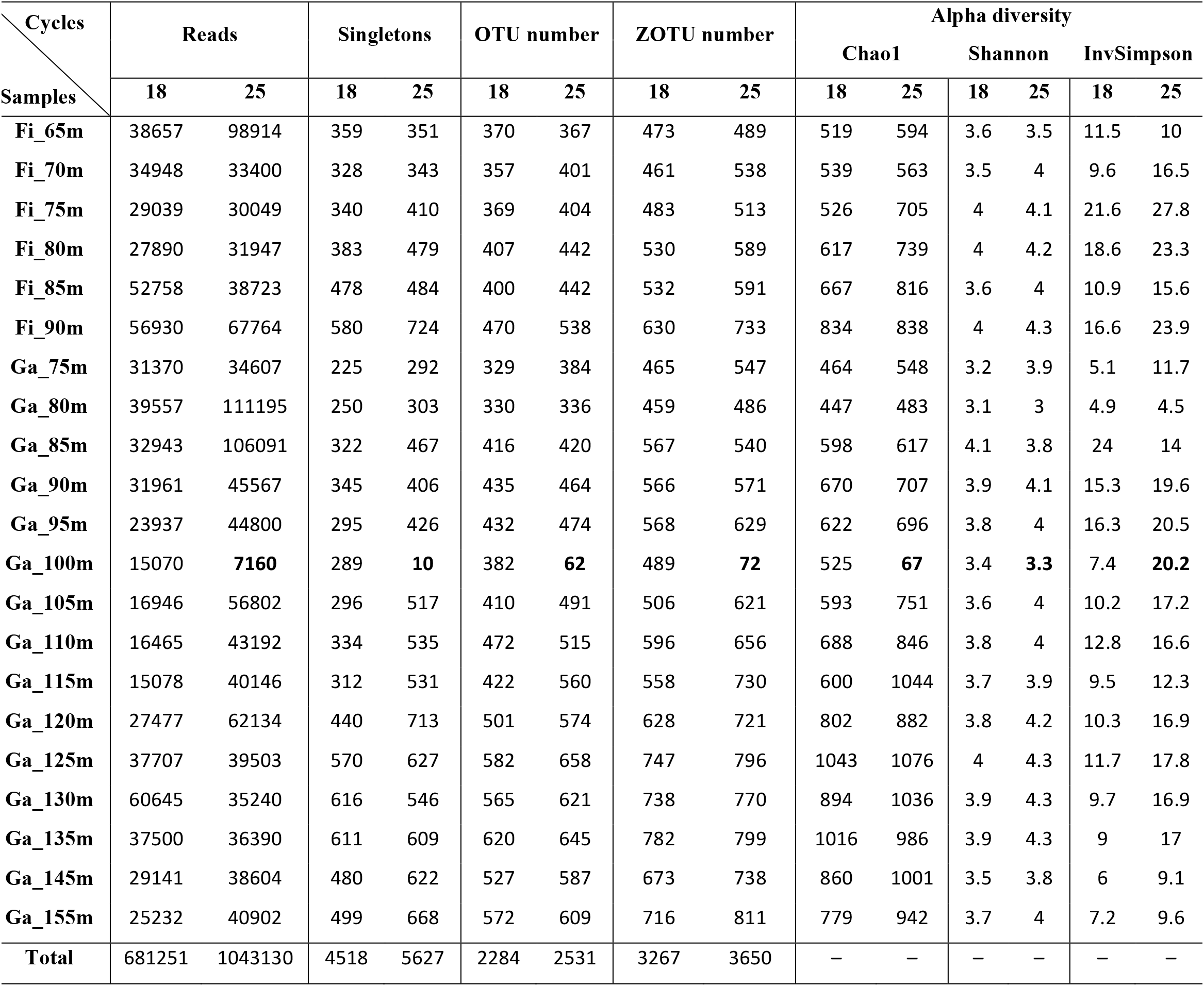
Comparison of the Lake Lugano 16S rRNA gene sequence data of two libraries prepared from the same original DNA, but with 18 and with 25 amplification cycles, respectively. The two libraries were sequenced together in a single Illumina run. Total reads and numbers of singletons (merged sequences occurring once in a sample) are based on raw reads. Total numbers of OTUs (97% sequence similarity), ZOTUs (zero-radius OTUs, i.e. amplicon sequence variants) and selected alpha diversity measures were determined after rarefying to even sequencing depths.

**Table S3.** ANOVA results showing the effects of experimental factors on different alpha diversity measures (Observed OTU richness, Chao1, Shannon and Inversed Simpson). The sequencing depth (i.e. read number per sample) had a significant effect (*) on the observed richness and Chao1, whereas the PCR amplification cycle number used for library preparation significantly affected Shannon and InvSimpson.

| Indices<br>Factors | Observed | Chao1 | Shannon | InvSimpson |
| --- | --- | --- | --- | --- |
| Sequencing<br>depth | F=11.68,<br>p=0.001 * | F=6.26,<br>p=0.017 * | F=0.04,<br>p=0.844 | F=0.463,<br>p=0.500 |
| Cycle number | F=1.36,<br>p=0.251 | F=1.79,<br>p=0.188 | F=8.46,<br>p=0.006 * | F=10.71,<br>p=0.002 * |

**Table S4.** t-Test results for differences in AOM rates (n = 3) in incubations prepared with North (NB) and South Basin (SB) water and amendments of different potential oxidants. After 16 and 32 days of incubations, rates were tested against the live control and the incubation with added sulfate. The significance level α set at 0.05 and p-values are given in parentheses. NS: not significant.

| Site | Treatment | 16 days |  | 32 days |  |
| --- | --- | --- | --- | --- | --- |
|  |  | Difference to control | Difference to sulfate | Difference to control | Difference to sulfate |
| NB | Nitrate | NS | > (0.05) | NS | > (0.005) |
|  | Nitrite | > (0.02) | > (0.01) | NS | NS |
|  | Molybdate | NS | NS | NS | NS |
|  | Sulfate | NS | – | NS | – |
|  | Control | – | – | – | – |
| SB | Nitrate | NS | NS | > (0.05) | NS |
|  | Nitrite | NS | NS | NS | NS |
|  | Molybdate | NS | NS | > (0.05) | NS |
|  | Sulfate | NS | – | NS | – |
|  | Control | – | – | – | – |

## Supplementary figures

**Figure S1.**
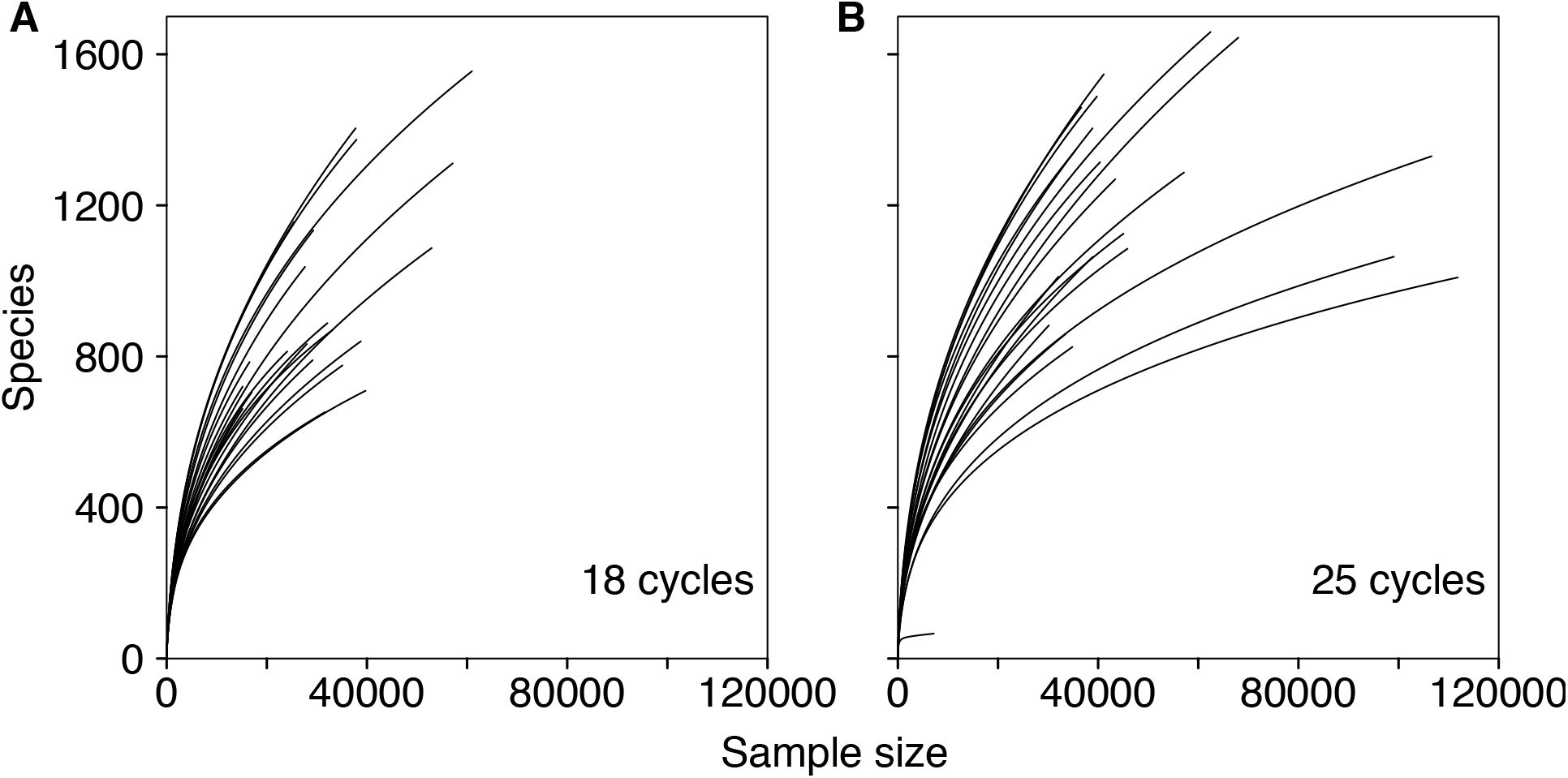
Rarefaction curves of samples from libraries generated with different PCR amplification cycles: (A) 18 cycles and (B) 25 cycles. Each curve represents one sample, showing the cumulative number of new OTUs (“Species”) in a given number of sampled sequences (“Sample size”).

**Figure S2.**
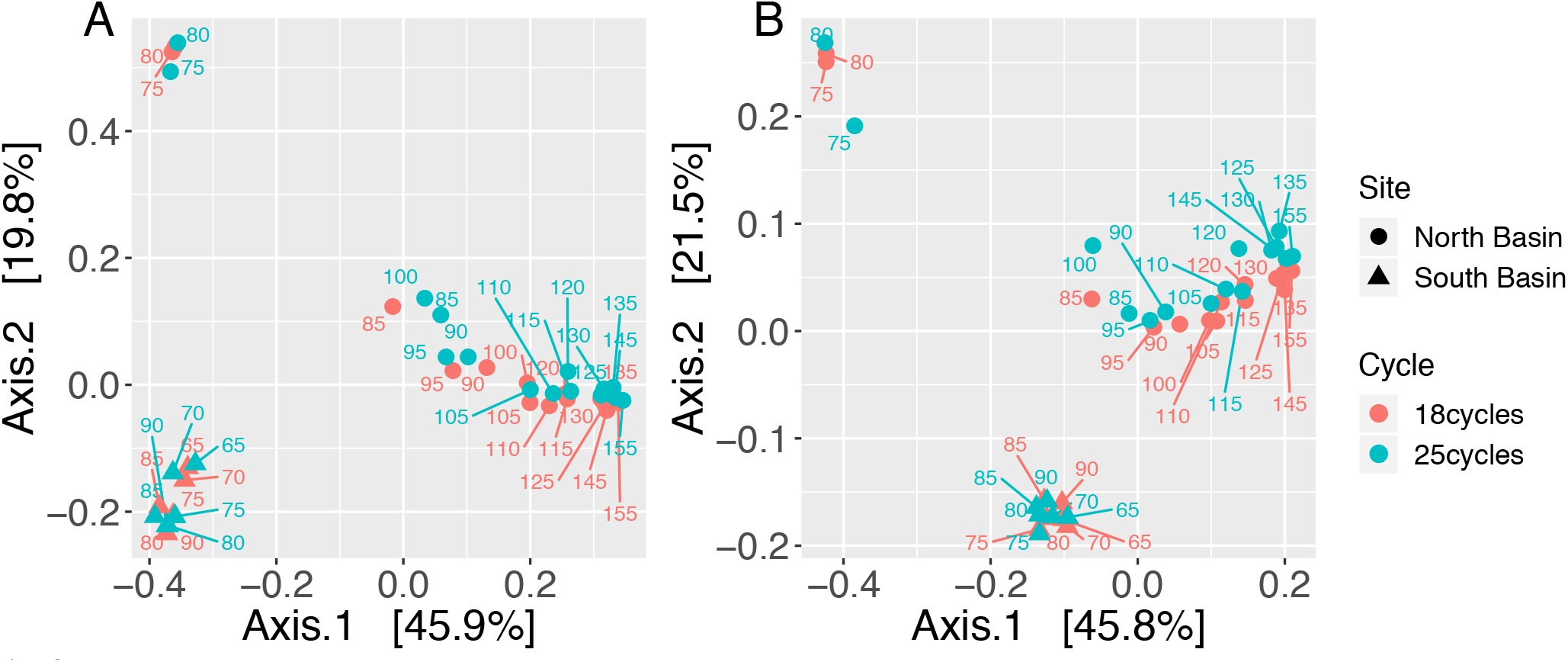
Principal coordinate analysis (PCoA) of microbial community structures in samples of the two basins (North Basin and South Basin) in Lake Lugano with two different PCR amplification cycle numbers (18 cycles and 25 cycles). Plots of PCoA on (A) Bray-Curtis and (B) weighted UniFrac distances revealed an atypical community structure for one of the samples (Ga_100 m, 25 cycles), possibly caused by a PCR amplification artifact. The x- and y-axes are indicated by the first and second coordinates, respectively, and the values in square brackets show the percentages of the community variation explained.

**Figure S3.**
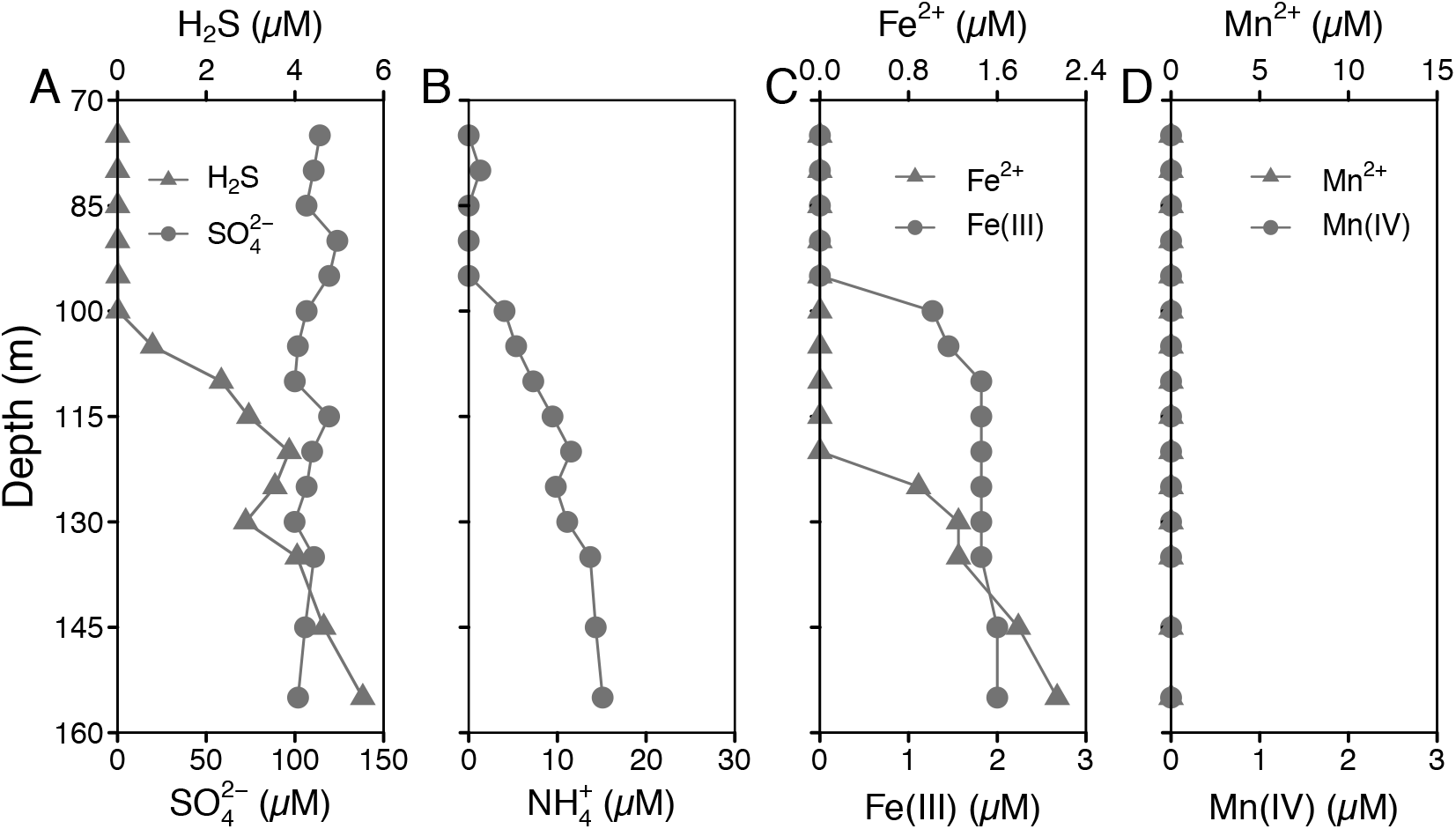
Concentration profiles of (A) sulfur species (sulfate and sulfide), (B) ammonium, (C) iron species and (D) manganese species in the water column of the permanently stratified North Basin of Lake Lugano in November 2016. Manganese species (both Mn^2+^ and Mn(IV) were below detection limit throughout the investigated depths).

**Figure S4.**
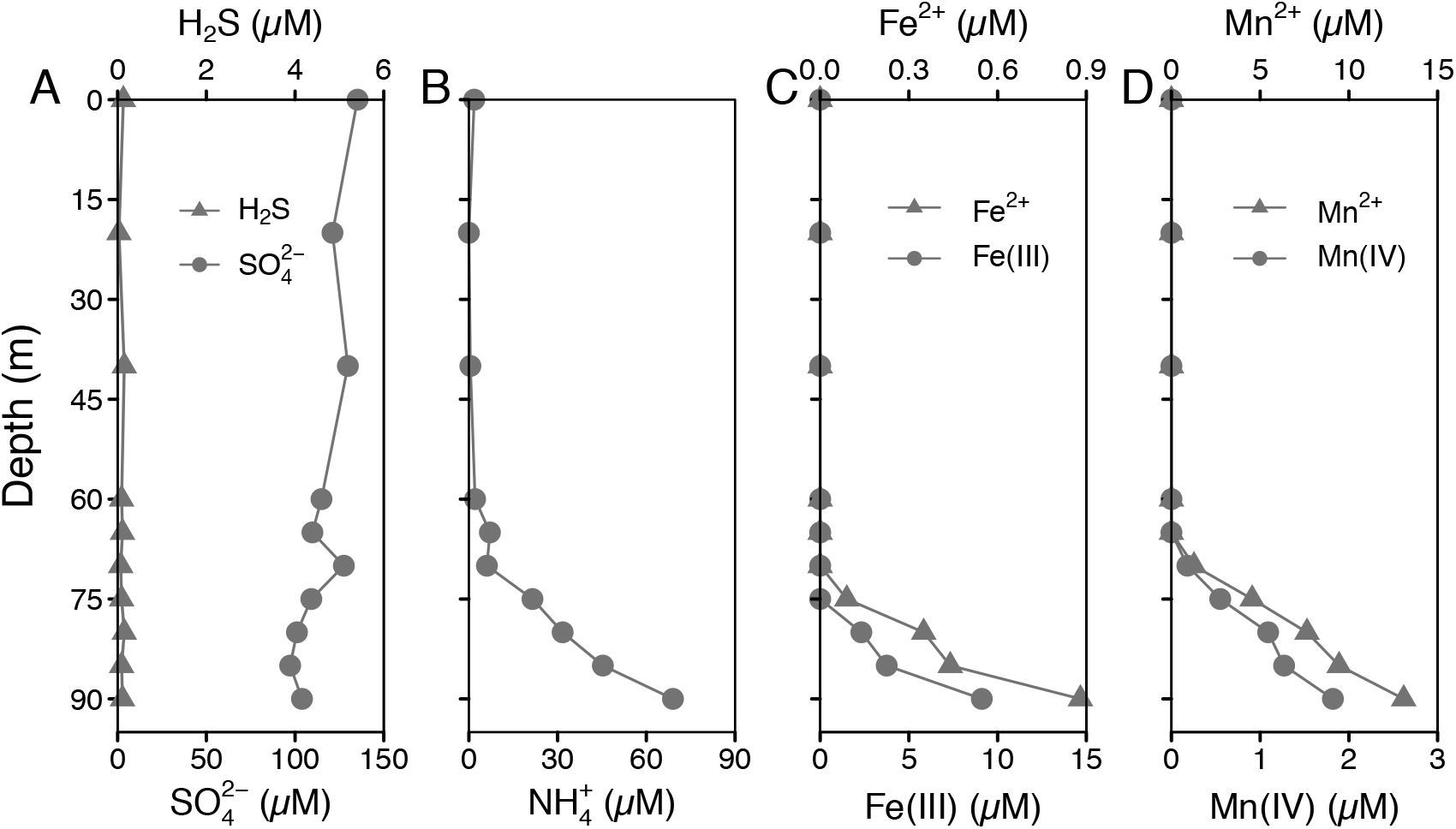
Concentration profiles of (A) sulfur species (sulfate and sulfide), (B) ammonium, (C) iron species and (D) manganese species in the water column of the seasonally stratified South Basin of Lake Lugano in November 2016.

**Figure S5.**
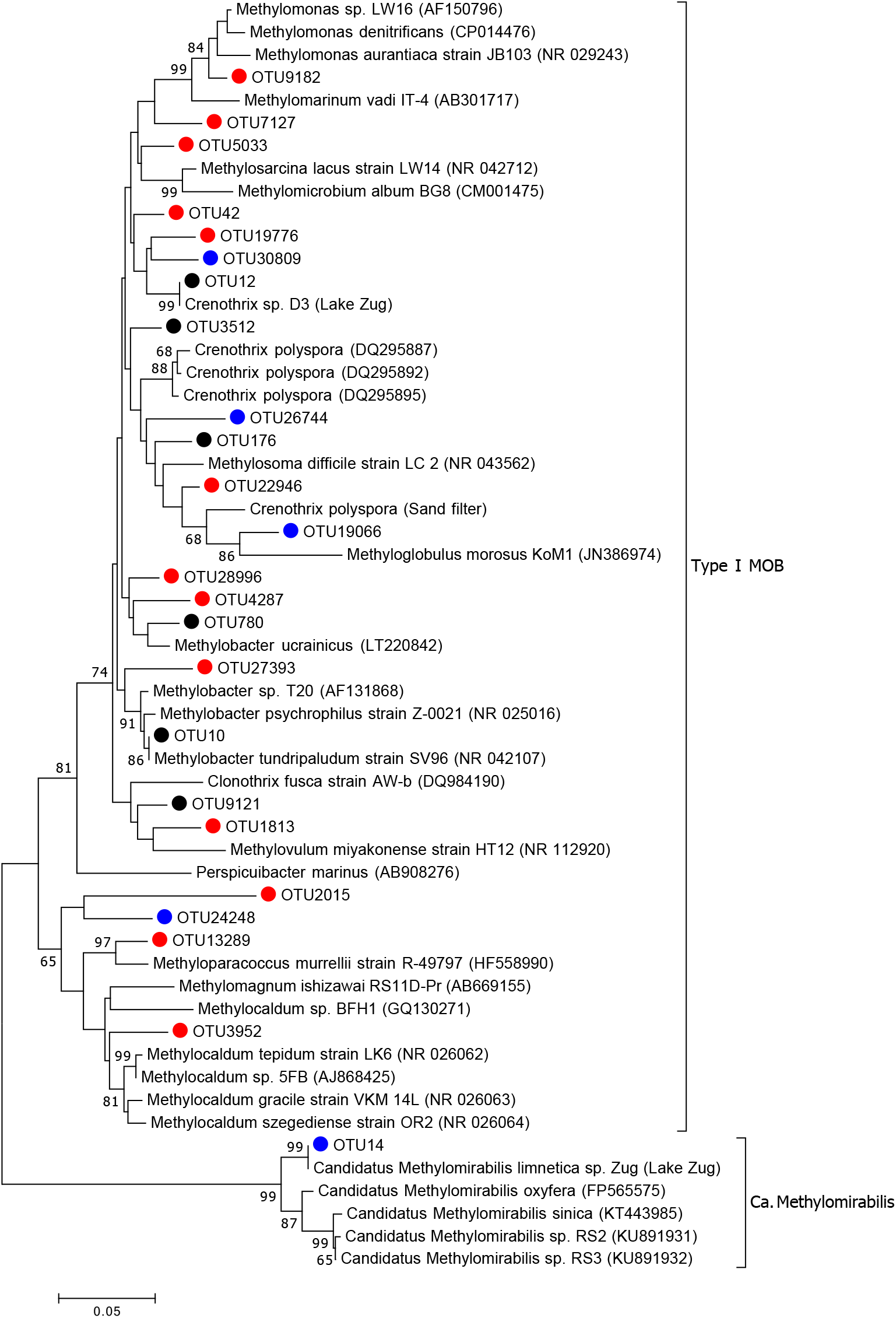
Neighbor-joining phylogenetic tree of 16S rRNA genes of both *Methylococcaceae* sp. and *Candidatus* Methylomirabilis detected in the South Basin (red bullets), in the North Basin (blue), and in both basins (black) where methane oxidation occurred. The tree was constructed using Maximum Composite Likelihood correction and partial gap deletion (Kumar et al. 2016), with a site coverage cutoff of 95%. Bootstrap values (> 65%) based on 1000 resampling are indicated at each node. Scale bar represents 5% of sequence divergence.

**Figure S6.**
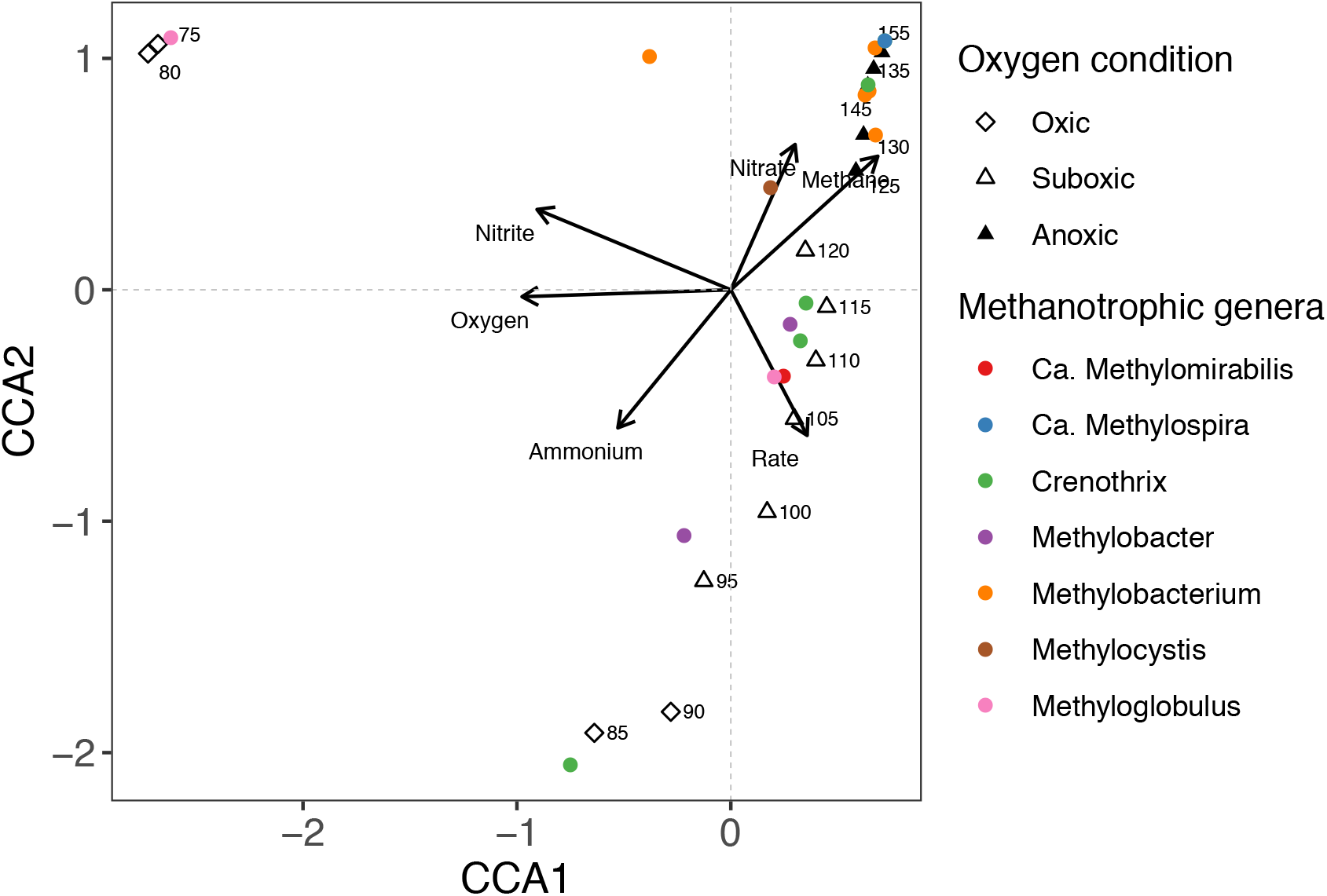
Canonical correspondence analysis (CCA) based on nutrient concentrations, AOM rate measurements, and potential methanotrophs detected in the North Basin of Lake Lugano. Triangles and diamonds represent samples in this basin under different oxygen conditions (with numbers indicating the water depths). Ordination was performed on the sequence data using Bray-Curtis distance. Taxa abundance and environmental variables (concentrations and rates) were Hellinger transformed prior to ordination. The CCA triplot shows that *Ca*. Methylomirabilis (red filled circle) is found in the suboxic water column of the permanently stratified North Basin (open triangles), and is positively correlated to the methane oxidation rate. Interestingly, the plot also shows that *Ca*. Methylomirabilis is anti-correlated to nitrite, suggesting that this taxon may be responsible for the depletion of the nitrite/nitrate pool in the habitat. Arrows represent solute concentrations, with arrowheads indicating their direction of increase.

**Figure S7.**
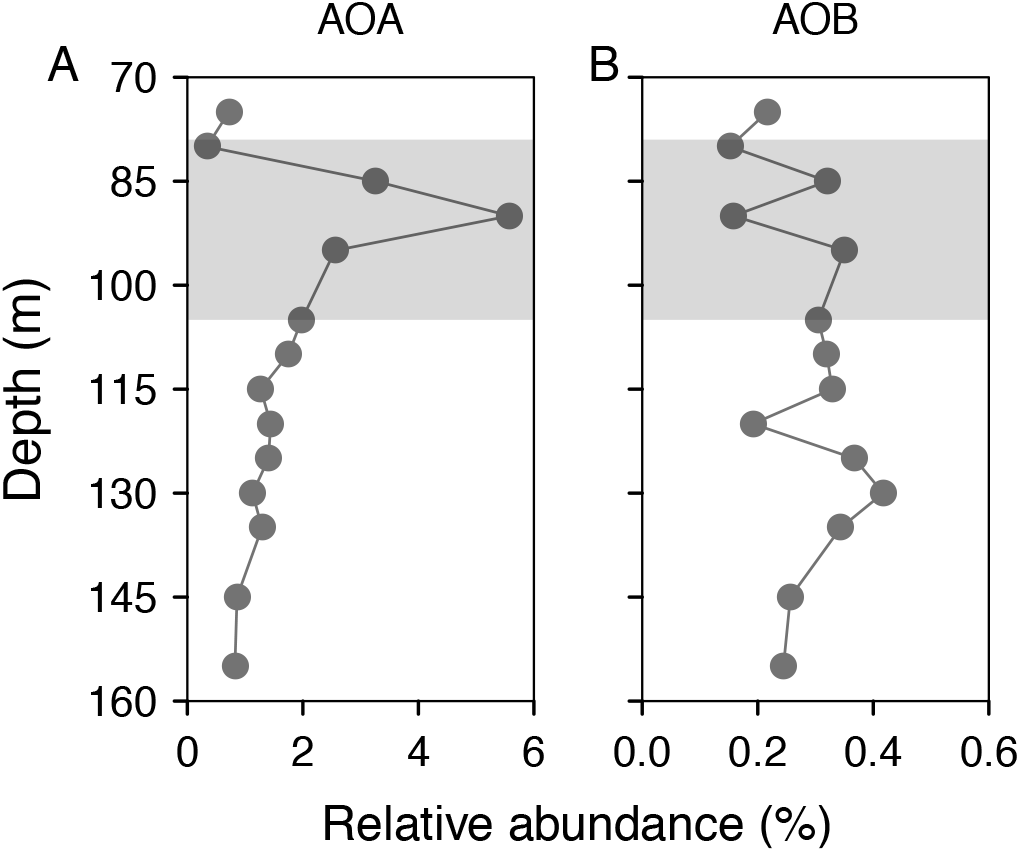
Depth profiles of relative abundances of (A) ammonium-oxidizing archaea (AOA, *Ca*. Nitrosopumilus and *Ca*. Nitrosoarchaeum), (B) ammonium-oxidizing bacteria (AOB, *Nitrosomonas* and *Nitrosospira*) across and below the redoxcline (indicated with grey) of the Lake Lugano North Basin in November 2016. Data are based on relative read abundances of 16S rRNA gene sequences.

**Figure S8.**
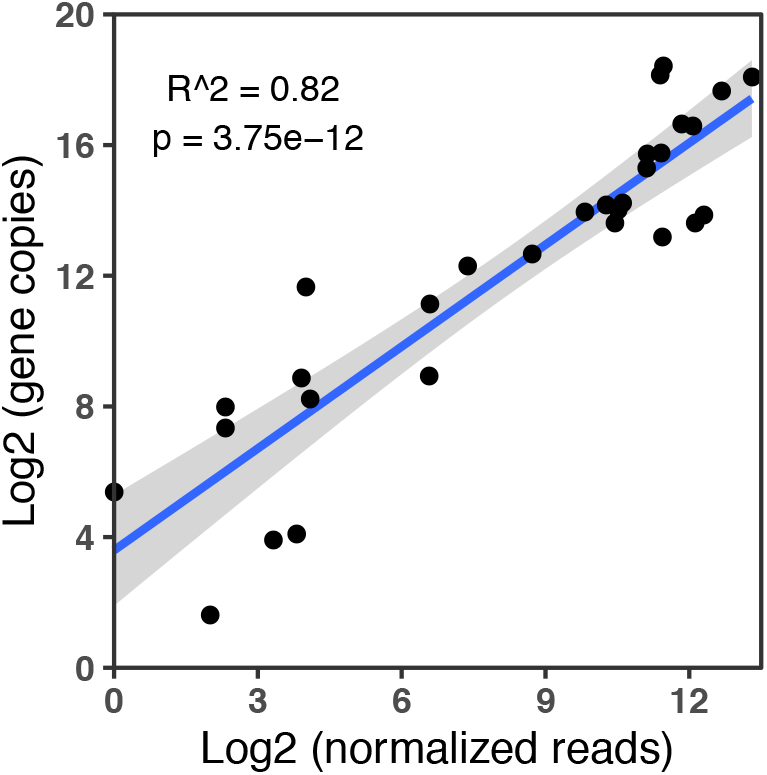
Relationship between 16S rRNA gene copy numbers and the normalized read numbers (to a same sequencing depth) of 16S rRNA gene sequences of *Ca*. Methylomirabilis in the North Basin water column. Samples are from October 2010, September 2014 and November 2016.

**Figure S9.**
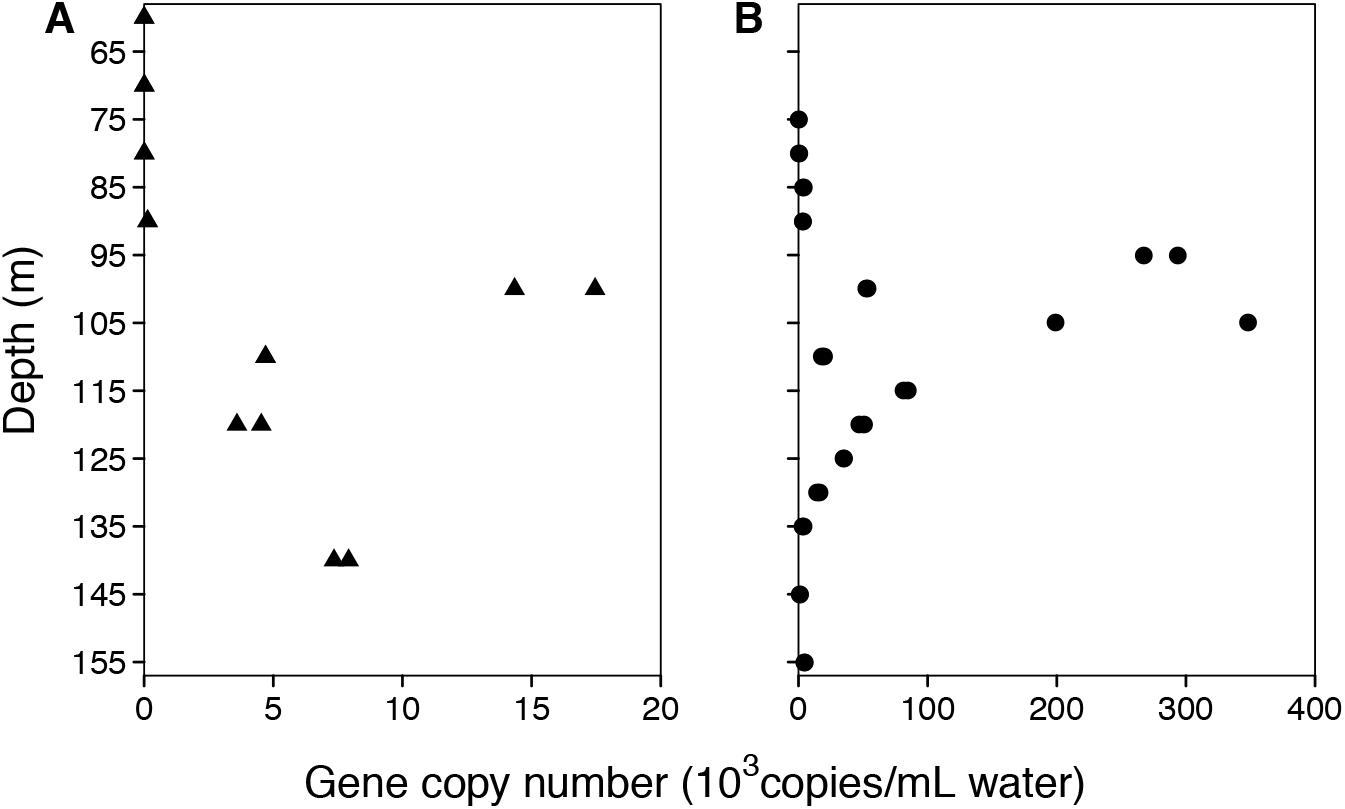
Depth profiles of 16S rRNA gene copy numbers of *Ca*. Methylomirabilis in the water column of the permanently stratified North Basin of Lake Lugano in (A) September 2014 and (B) November 2016. The redoxcline was located between 104 and 125 m in 2014, and between 79 and 105 m in 2016. Based on qPCR data (i.e., the local maximum gene copy numbers between the two years), the estimated apparent doubling time of *Ca*. Methylomirabilis was approximately 6 months, longer than the anoxic stratification period of ∼5 months in the South Basin.

